# Testing for batch effect through age predictors

**DOI:** 10.1101/531863

**Authors:** Polina Mamoshina, Kirill Kochetov, Evgeny Putin, Alex Aliper, Alex Zhavoronkov

## Abstract

Transcriptome profiling has been shown really useful in the understanding of the aging process. To date, transcriptomic data is the second most abundant omics data type following genomics. To deconvolute the relationship between transcriptomic changes and aging one needs to conduct an analysis on the comprehensive dataset. At the same time, biological aging clocks constructed for clinical use needs to robustly predict new data without any further retraining. In this paper, we develop a transcriptomic deep-learned age predictor. Deep neural networks (DNN) are trained and tested on more than 6 000 blood gene expression samples from 17 datasets. We apply methods based on output derivatives of DNN to rank input genes by their importance in age prediction and reduce the dimensional of the data. We also show that batch effect in transcriptome datasets of healthy humans is indeed significant, but the existing normalization techniques, while removing technical variation quite effectively, also remove age-associated changes. So robust methods of age prediction are needed.

## Introduction

The rates of aging may vary substantially among the different individuals and population groups and are significantly influenced by the environmental and hereditary factors. Multiple attempts have been made to develop the biologically-relevant biomarkers of human aging.

However, the biomarkers proposed so far usually focus on monitoring a restricted number of processes known for being directly correlated with the chronological age such as the telomere length-based or DNA methylation. There is a need for the biologically-relevant quantifiable, interpretable and therapeutically-targetable multi-modal biomarkers of aging. Even though these clocks were developed using traditional machine learning approaches as a linear regression with regularization the results suggest that gradual changes during aging can be tracked using various data types with reasonable accuracy.

Previous studies demonstrated age-associated changes in the transcriptome of model organisms [1] and multiple human tissues [2, 3, 4]. In 2015, Peters and colleagues performed the massive analysis of transcriptional profiles of aging and used six blood expression profiles (7,074 samples in total) to build a predictor of age with leaving a dataset out as validation [3]. Using elastic-net regularized linear regression, their approach achieved an average MAE of 7.8 years. In the analysis, 1,497 genes were identified as age-related. In 2018, Mamoshina et al proposed a panel of transcriptomic age predictors and the approach of comparing different methods of selecting age-related genes [4]. A deep neural network was the most accurate age predictor showing the accuracy of 0.91 in terms of Pearson correlation and mean absolute error of 6.14 years. Further validation on the external GTEx dataset showed the accuracy of 0.80 with respect to the actual age bin prediction. Another promising finding was that the list of the features most relevant to age prediction identified by the deep neural network is the closest results to the final consensus ranking produced by other ML age predictors suggesting the superior generalization abilities.

However, most the age predictors so far use a limited number of samples from a relatively small cohort of people for independent validation. The impact of the technical variability or so-called batch effect on the age prediction also remains largely unaddressed in the literature. This remains a key challenge in developing of aging biomarkers that can be used in the clinical setting as they should robustly predict the age of previously unseen samples from an independent dataset.

In this work, we decided to use deep learning models for predicting age of humans by their gene expression profiles as they demonstrated impressive results on blood biochemistry and cell counts [5, 6, 7], transcriptomics [4], microbiome [8], facial images [9], bone X-ray images [10], brain MRI images [11].

Here we firstly collect a large dataset (6465 samples from 17 datasets) of transcriptomic datasets of healthy and diseased human blood samples profiled using two microarray platforms (Illumina HT12 v3.0 and v4.0). We then identify the technical variability in blood transcriptome and showing that has a stronger impact on expression than disease state or age. We apply several normalization techniques, showing that while some of them are quite effective in removing the batch effect, they also mitigate age-associated changes in the blood transcriptome.

## Materials and Methods

### Data

Gene expression profiles were collected from the publicly available repository Gene Expression Omnibus (https://www.ncbi.nlm.nih.gov/geo/). In total, we analyzed 6465 transcriptomic samples, labeled according to the chronological age of the tissue samples’ donors, from 17 datasets (Table 1).

**Table 1.** List of datasets analysed with number of samples. Nine expression datasets were selected as training set and eight were used as external testing set. Only healthy samples were selected

| GEO Accession | N of samples | Tissue | N of selected samples |
| --- | --- | --- | --- |
| *Training* |  |  |  |
| GSE65907 | 2112 | PBMC | 2112 |
| GSE56045 | 1202 | Monocytes | 1202 |
| GSE33828 | 881 | Whole blood | 881 |
| GSE78840 | 653 | Whole blood, Lymphocytes | 653 |
| GSE61672 | 546 | Whole blood | 126 |
| GSE35846 | 128 | Whole blood | 128 |
| GSE63060 | 326 | Whole blood | 104 |
| GSE47728 | 228 | Whole blood | 228 |
| GSE47727 | 122 | Whole blood | 122 |
| Total for training |  |  | 5555 |
| *Testing* |  |  |  |
| GSE102008 | 733 | Whole blood | 94 |
| GSE107990 | 671 | PBMC | 171 |
| GSE63061 | 388 | Whole blood | 134 |
| GSE86434 | 249 | Lymphocytes, Whole blood | 249 |
| GSE56580 | 214 | Lymphocytes | 214 |
| GSE94496 | 208 | Monocytes | 208 |
| GSE74629 | 50 | Whole blood | 14 |
| GSE75025 | 24 | PBMC | 24 |
| Total for testing |  |  | 910 |
| **Total** |  |  | **6465** |

### Train and test set design

To evaluate the performance of the age predictors robustly, we selected three datasets as external testing sets. We also selected only samples of healthy subjects for the training purposes, for age predictors to fit to age-associated changes in the transcriptome, rather than disease-associated changes.

### Normalization techniques

We compare non-normalized data (raw) to the following commonly used for gene expression normalization methods:

1. Normalization by Reference Gene (RefGenNorm) calculated as following:

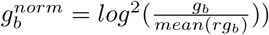

where *g* is the gene to normalize, *rg* is the reference gene (we choose *C*1*ORF* 43), *b* is a some batch (GEO in our case). Reference genes or housekeeping genes should have the constant level of the expression across different tissues and experimental designs, diseases. Here we explored the Chromosome 1 Open Reading Frame 43 (C1orf43) gene as a reference one (Figure 1), that has been shown to have the most stable expression in different human tissue and experiments[12].
2. Cross-platform normalization method (XPN) proposed by Shabalin et al[13].
3. Quantile normalization (QN) [14]
4. Distribution Transformation (DisTran) [15]

**Figure 1.**
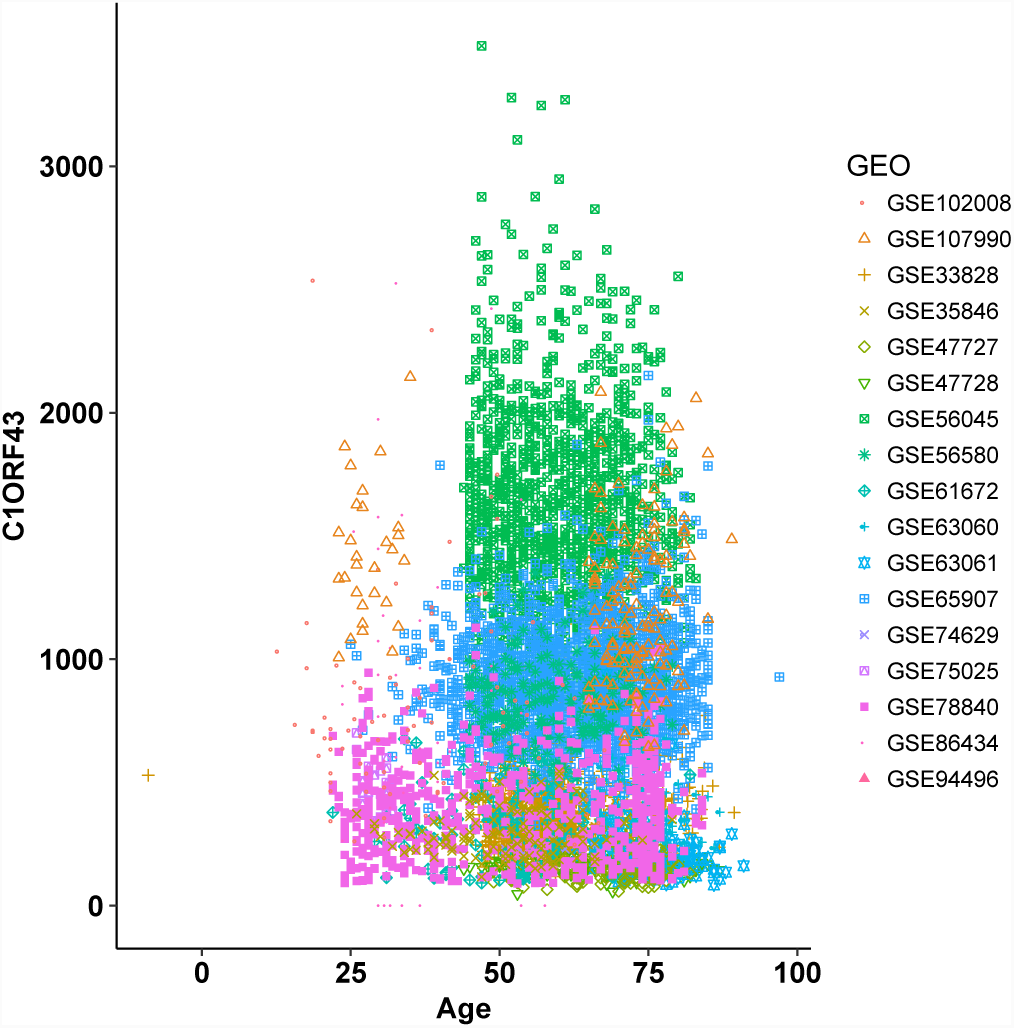
Distribution of the Chromosome 1 Open Reading Frame 43 (C1orf43) gene expression across datasets.

In contrast to RefGenNorm method, which is independent of other datasets, the latter three methods require the reference dataset for data transformation. Here we have selected GSE65907 as the most comprehensive one. We used R implementation of XPN, QN and DisTran normalizations developed by Rudy et al [16].

Despite applying normalization provided above, there are a lot of outliers left in the normalized data. To handle it we used the standard technique for outliers detection and exclusion called three-sigma rule [17]. We adapted this rule for our large dimensional data (13454 genes) and classified sample as an outlier if it did not satisfy the three-sigma rule over more than 99% of genes within each of four presented tissues. Without such an adaptation we would lose more than half of the data.

### Regression model implementation

We adapted a deep feed-forward neural network with a weighted layer providing a way to rank the input features and ElasticNet-based regularization to this weighted layer providing the ability to control sparsity/smoothness of weights[18]. We build and train models using Keras (https://keras.io) library with Tensorflow (https://www.tensorflow.org/) backend. Grid search over a space of model parameters with five-fold cross-validation was used in order to find the best performing neural network architecture. The best model was the one trained on 1000 selected genes (by deep feature selection method) and has four layers with accordingly 800, 700, 600 and 500 neurons, Exponential Linear Unit (ELU)[19] activation function after each hidden layer. Also, we used Adam[20] algorithm for loss function optimization. For the purposes of regularization, we used dropout[21] with 15% probability after each layer and combined ridge and lasso regularization with 10*e*^−6^ coefficients as additional loss term.

All experiments were conducted using an NVIDIA 1080Ti (Pascal) graphics processing unit.

### Model evaluation

We used the following metrics to evaluate the accuracy of age prediction models:

1. Mean absolute error: 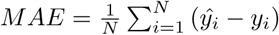
where *ŷ*_*i*_ is a predicted age of a sample *i, y*_*i*_ is the chronological age value of a sample *i*, and N is a number of samples. *MAE* demonstrates average disagreement between the chronological age and the predicted age. *MAE* of 0 means that the predicted age and actual age are in a perfect agreement.
2. Coefficient of determination:

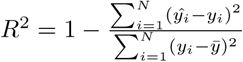

where *y*_*i*_ is the chronological age value of a sample *i, ŷ*_*i*_ is the predicted age value of a sample *i*, and 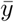 the mean chronological age in the distribution. *R*^2^ shows the percentage of variance explained by the regression between predicted and actual age. *R*^2^ of 1 demonstrates that all of the variance in the actual age explained by the prediction.

## Results

To examine associations between transcriptional changes in blood and chronological age, we collected and analyzed 17 datasets. We collected 6465 samples of 13454 gene expression values. The mean age of the collected samples was 61 years (Figure 2).

**Figure 2.**
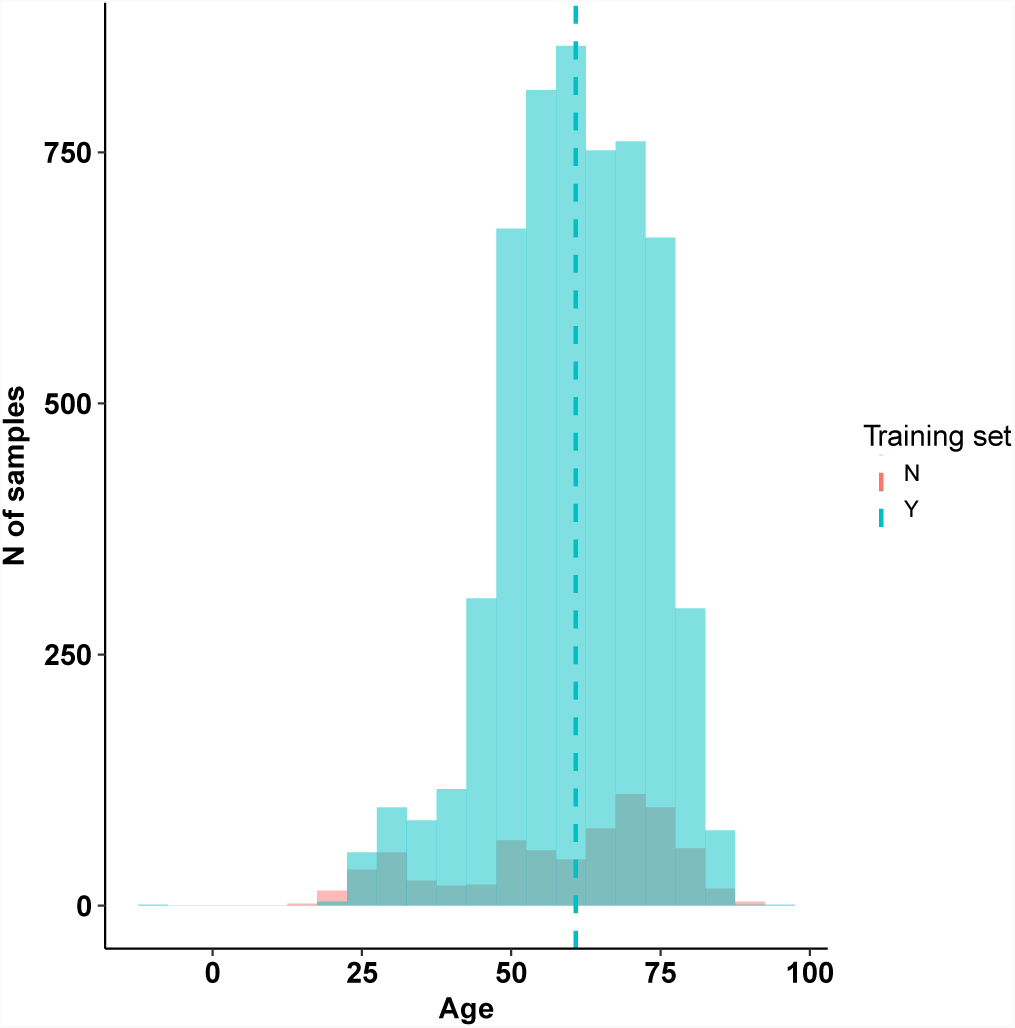
Age distribution of samples in the training and testing sets.

To distinguish batch effect from biological differences between different samples, we performed t-Distributed Stochastic Neighbor Embedding (t-SNE) [22] and visualized first two components of the manifold (Figure 3). We compare several methods of adjusting for batch effect. Compared to other methods, the XPN algorithm demonstrated great improvement in batch effect removal, improving the entanglement of datasets (Figure 3).

**Figure 3.**
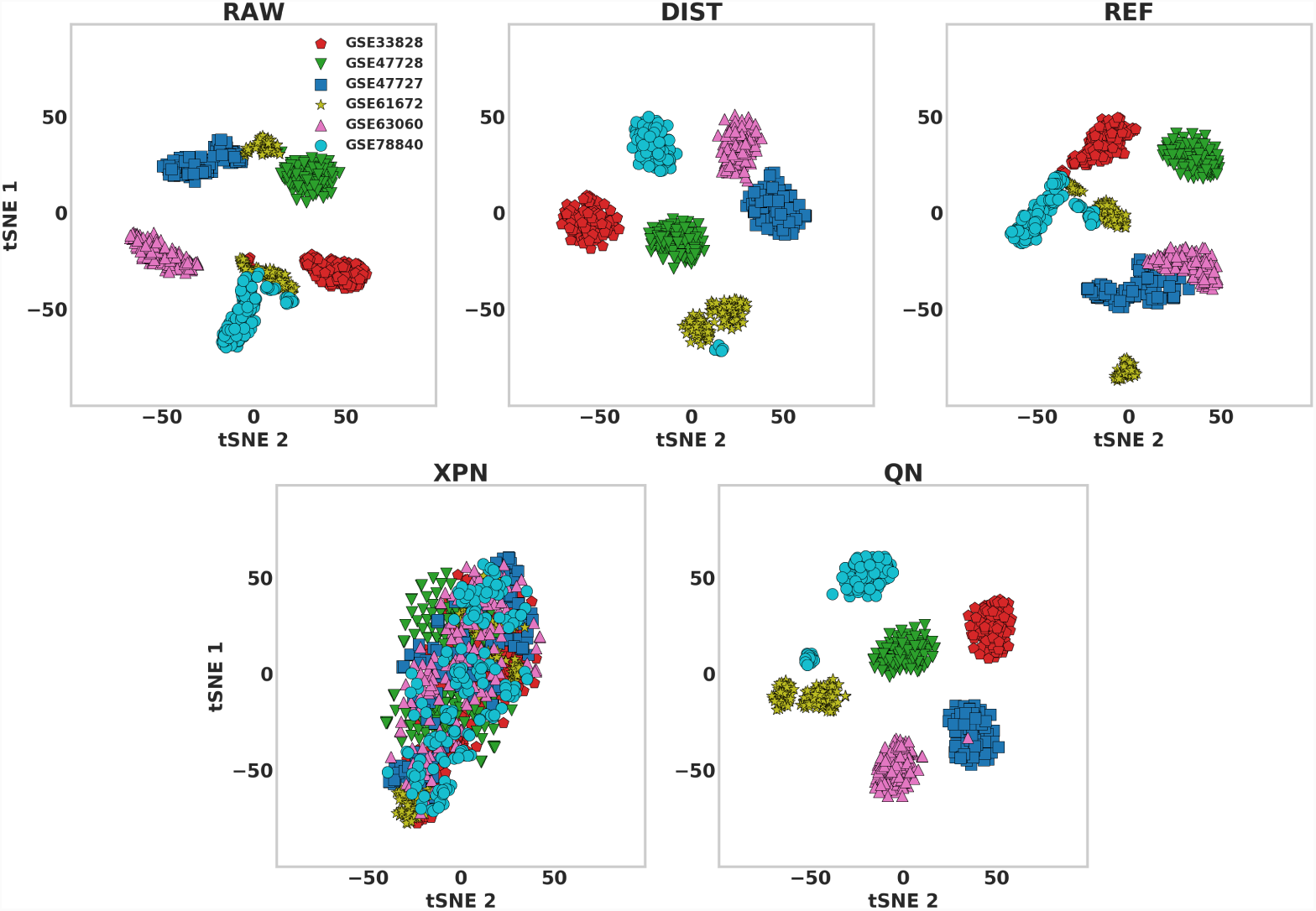
Visualization of presence of batch effect in raw and normalized datasets. RAW is for raw data, DIST is for data normalized with Distribution Transformation. REF is for normalization by Reference Gene. XPN is for cross-platform normalization method proposed by Shabalin et al[13]. QN is for quantile normalization.

Age prediction results are summarized in Table 2 and showed in Figure 4. Baseline accuracy, where all samples are predicted as the median for distribution, is 8.14 years in terms of *MAE* and *R*^2^ of 0.50 on the testing set and 9.07 *MAE* and *R*^2^ of 0.12 on the training set. DisTran method achieved the highest accuracy on cross-validation compared to other methods (*MAE* of 5.05 years and *R*^2^ of 0.70) and the accuracy lower than the baseline on testing set (*MAE* of 9.29 and *R*^2^ of 0.44). Quantile normalization showed relatively good accuracy on cross-validation (*MAE* of 5.41 years and *R*^2^ of 0.66), which was reduced, however, on the testing set to accuracy compared to baseline accuracy. At the same time, RefGenNorm normalization displayed the greater improvement compared to baseline, with *MAE* of 7.98 years and *R*^2^ of 0.61. It is interesting because only RefGenNorm transformation of the raw data is truly independent of other datasets. While XPN algorithm demonstrated the best results in mitigation of batch effect, the model trained on this data only achieved 15.68 years *MAE* and *R*^2^ < 0 on the testing set (model fits worse than a median).

**Table 2.** The performance of models trained on the five data before (Raw) and after cross-dataset normalization (RefGenNorm, XPN, QN, DisTran, See Methods for details).

| Data | $MAE$ (years) ↓ | $R^2$ ↑ |
| --- | --- | --- |
| Baseline | 9.07 | 0.12 |
|  | 8.14 | 0.50 |
| Raw | 5.42 | 0.65 |
|  | 9.15 | 0.52 |
| RefGenNorm | 5.99 | 0.58 |
|  | **7.98** | **0.61** |
| XPN | 5.77 | 0.61 |
|  | 15.68 | <0.00 |
| QN | 5.41 | 0.66 |
|  | 8.19 | 0.55 |
| DisTran | **5.05** | **0.70** |
|  | 9.29 | 0.44 |

**Figure 4.**
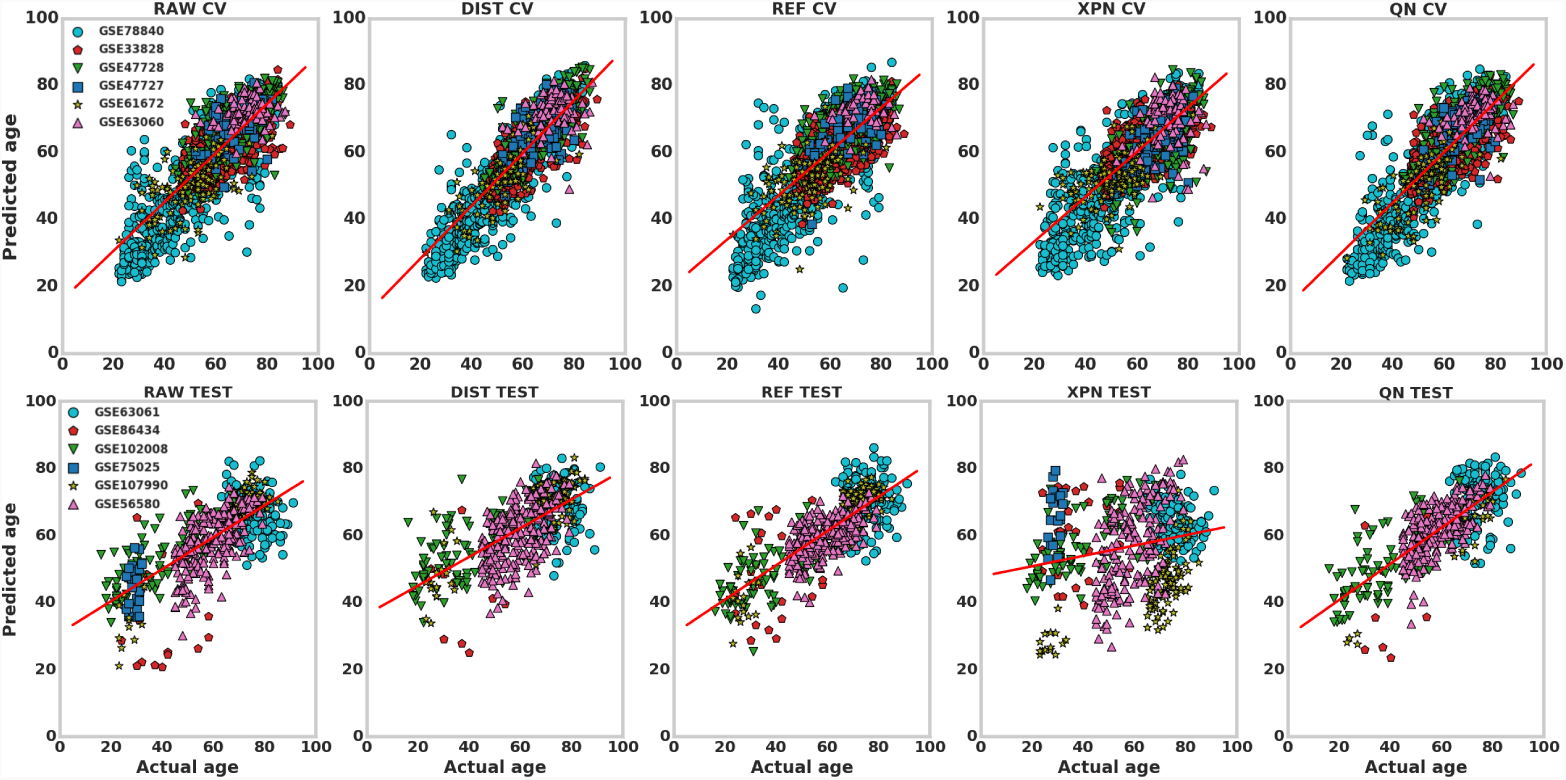
Actual vs. predicted age plots on cross-validation and testing sets before and after normalization. RAW is for raw data, DIST is for data normalized with Distribution Transformation. REF is for normalization by Reference Gene. XPN is for cross-platform normalization method proposed by Shabalin et al[13]. QN is for quantile normalization.

We compared the batch effect for the top 10, 50, 100, 500 and 1000 the most important genes for the age prediction identified by the model (Figure 5). Interestingly, top-ranked genes (up to 20) are not related to batch, suggesting that DNNs still capture genes that are strongly related to age rather than batch effect. It is critical because age distributions of different datasets are indeed different and could bring certain bias.

**Figure 5.**
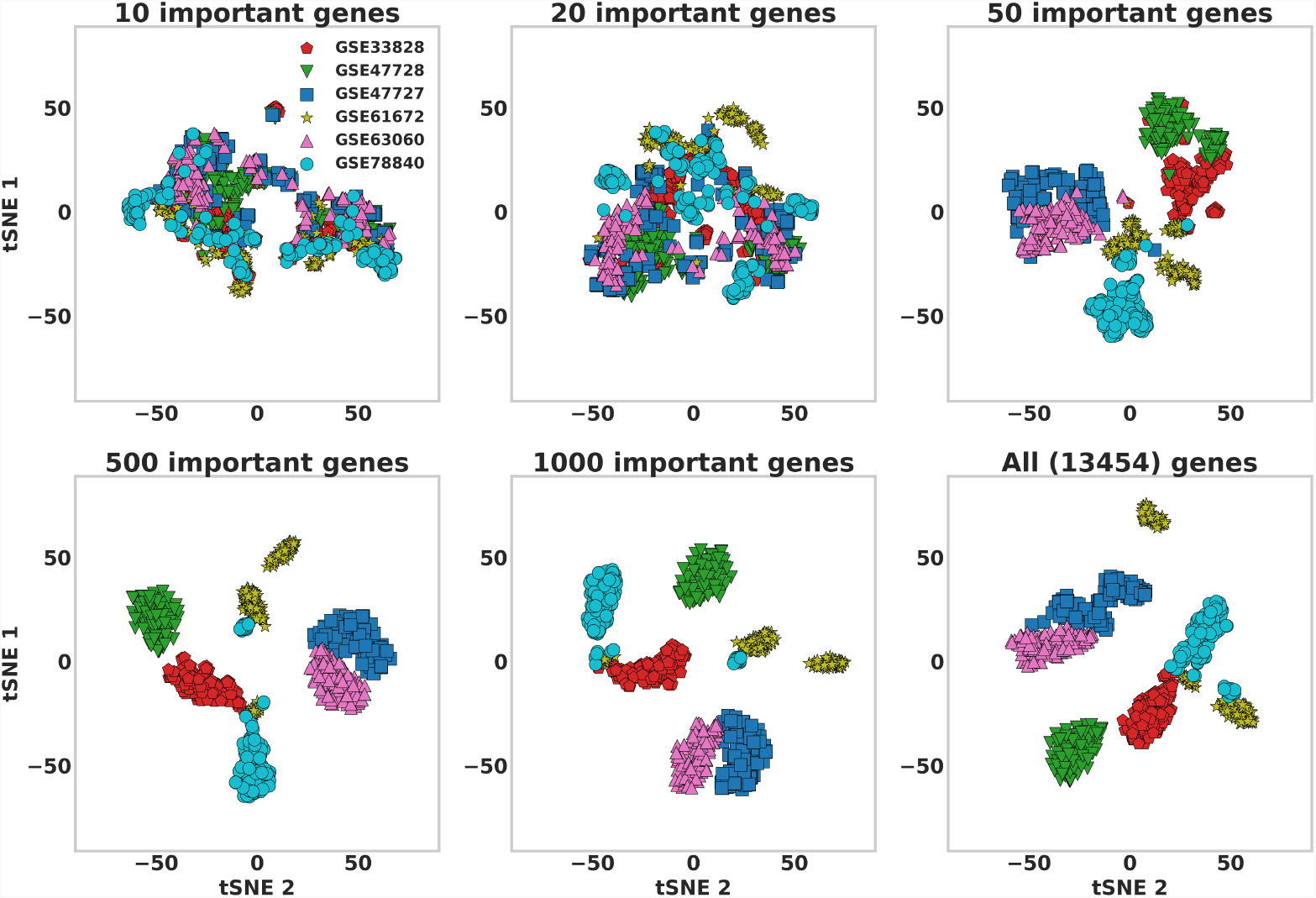
Visualization of presence of batch effect using different amount of important genes selected by DFS

## Discussion

In this article, we illustrated the size of the batch effect in 17 publicly available gene expression datasets of healthy human blood. We compared several standard normalization methods with non-transformed data, showing that normalization by reference gene returning the best results in terms of accuracy of age prediction. We also showed that while some methods are removing batch effect significantly, also removing age-related changes. Given the magnitude of batch effect in transcriptomic datasets and that, cross-validation cannot replace independent-data validation for transcriptomic aging biomarkers.

Effective batch-effect removal techniques remains a key challenge for transcriptomic aging markers.

